# Cell-Surface LAMP1 is a Senescence Marker in Aging and Idiopathic Pulmonary Fibrosis

**DOI:** 10.1101/2025.03.04.640878

**Authors:** Gabriel Meca-Laguna, Michael Qiu, Anna Barkovskaya, Apoorva Shankar, Michael Rae, Amit Sharma

## Abstract

The accumulation of senescent cells (SEN) with aging produces a chronic inflammatory state that accelerates age-related diseases. Eliminating SEN has been shown to delay, prevent, and in some cases reverse aging in animal disease models and extend lifespan. There is thus an unmet clinical need to identify and target SEN while sparing healthy cells. Here, we show that Lysosomal-Associated Membrane Protein 1 (LAMP1) is a membrane-specific biomarker of cellular senescence. We have validated selective LAMP1 upregulation in SEN in human and mouse cells. Lamp1^+^ cells express high levels of prototypical senescence markers p16, p21, Glb1, and have low Lmnb1 expression as compared to Lamp1^-^ cells. The percentage of Lamp1^+^ cells is increased in mice with fibrotic lungs after bleomycin instillation. Finally, we use a dual antibody-drug conjugate (ADC) strategy to eliminate LAMP1^+^ senescent cells.

## Introduction

Throughout life, cells are continually exposed to a myriad of different stressors which include oncogenic stress, telomeric shortening, wounds, infections, irradiation, certain drugs, etc. If the cellular insult constitutes irreparable damage, programmed cell death pathways provoke apoptosis in the affected cells. However, most such challenges can be resolved, in which case damaged cells arrest their cycle in order to eventually restore homeostatic conditions. Cellular senescence, initially described in 1961 by Leonard Hayflick (Hayflick & Moorhead, 1961), is an irreversible cell cycle arrest following cellular insult (Munoz-Espin & Serrano, 2014). Phenotypically, senescent cells (SEN) secrete a cocktail of pro-inflammatory and other molecules known as the senescence-associated secretory phenotype (SASP), which contributes to the chronic, sterile, low-grade inflammation of aging (inflammaging), accelerating morbidity onset (Munoz-Espin & Serrano, 2014; Xu et al., 2018). Accordingly, transplantation of senescent cells induces physical dysfunction in mice (Xu et al., 2018). Conversely, eliminating senescent cells increases health- and lifespan (Kirkland & Tchkonia, 2017). Thus, senescent cells contribute to age-related decline. For this reason, there is a growing interest in developing interventions that eliminate SEN or modulate their inflammatory secretory phenotype (Kirkland & Tchkonia, 2017).

So far, the combination of Dasatinib and Quercetin (D+Q), Navitoclax, and Fisetin are the most widely tested senolytics, which target apoptotic vulnerabilities in SEN (Chang et al., 2016; Yousefzadeh et al., 2018; Zhu et al., 2015). Additionally, the immune system executes surveillance on senescent cells (Kale et al., 2020), and investigators have exploited chimeric antigen receptor (CAR) biotechnology to target SEN with engineered lymphocytes, leveraging surface proteins like uPAR or stress-associated NKG2D ligands (Amor et al., 2020; Deng et al., 2024; Yang et al., 2023). Small molecule approaches show promising results *in vitro* and *in vivo* in models, but limited efficacy has been observed in human clinical trials (Nambiar et al., 2023) or in genetically heterogeneous mice in the Interventions Testing Program (ITP) (Harrison et al., 2024). Other strategies that have been used to target SEN *in vitro* and *in vivo* on the basis of their surface proteome (surfaceome) include antibody-drug conjugates (ADCs) against B2M and ApoD (Poblocka et al., 2021; Takaya et al., 2023) and antibody-dependent cell-mediated cytotoxicity (ADCC) against DPP4 (Kim et al., 2017).

Despite the well-documented role of SEN in aging and age-related diseases, the lack of a robust biomarker for characterizing senescence in tissue samples is a significant impediment in the field. Such a biomarker should be agnostic to the tissue and cell type of origin and the senescence-causing insult. The lack of such a biomarker limits researchers’ ability to evaluate SEN elimination (senolysis) after candidate senolytic treatments.

One of the most well-documented hallmarks of SEN is their increased lysosomal content and activity (Dimri et al., 1995; Hernandez-Segura et al., 2017; Rovira et al., 2022). This hallmark feeds into multiple other features of senescence, such as changes in morphology, the SASP phenotype, and metabolic alterations. Indeed, early efforts quantifying lysosomal activity resulted in the discovery of senescence-associated β-galactosidase (SA-β-Gal), one of the most widely used biomarkers of senescence (Dimri et al., 1995).

Lysosomal Associated Membrane Protein 1 (LAMP1, also known as CD107a) is a master orchestrator of the structural integrity of lysosomes (Eskelinen, 2006). It is estimated that LAMP1 (and LAMP2) constitute half of all proteins of the lysosomal membrane (Eskelinen, 2006; Hunziker & Geuze, 1996). LAMP1, as a type I transmembrane glycoprotein, is mostly localized in late endosomes and lysosomes (Hunziker & Geuze, 1996), and is a heavily N-glycosylated protein (Alessandrini et al., 2017). In immune cells, CD107a is also a cell-surface marker of immune activation and cytotoxic degranulation (Alessandrini et al., 2017; Krzewski et al., 2013); however, it can only be detected using inhibitors of intracellular protein transport (such as GolgiStop, monesin-containing, or brefeldin A-containing reagents) since its expression on the membrane is transient and the protein is internalized rapidly. Therefore, LAMP1 is only briefly found at the cell surface of healthy cells due to the fusion of lysosomes with the plasma membrane, and thus, mostly undetectable. In the context of disease, it has been reported that melanoma, carcinoma, and fibrosarcoma cells exhibit abnormal localization of LAMP1 on their plasma membranes, possibly as part of repair of the damaged cell surface (Corrotte et al., 2015; Reddy et al., 2001; Sarafian et al., 1998). Cell-surface LAMP1 has been associated with more malignant and metastatic cancers, where it promotes drug resistance through increased lysosomal exocytosis (Alessandrini et al., 2017).

In summary, the ability to identify and characterize senescent cells is crucial for understanding their role in aging and developing targeted interventions. Here, we describe LAMP1 as a cell surface-specific marker of senescence. LAMP1’s presence on the cellular membrane is highly increased in human and mouse SEN. In mouse tissue, cells expressing Lamp1 on their surface showed features of senescence. Additionally, senescence induction in the lungs of mice using bleomycin caused an increase in Lamp1^+^ cells. Finally, SEN are eliminated using a LAMP1-targeting antibody. These findings describe a biomarker that can be leveraged to further understand and target senescent cells.

## Results

### Multi-omic screens identify LAMP1 upregulation in cellular senescence and co-expression with p21

We analyzed a publicly available proteomic screen of SEN plasma membrane to identify proteins specifically enriched on the surface of senescent cells (Marin et al., 2023). This screen contains proteins enriched on SEN of several cell types of origin and modes of senescence induction (**Figure 1a**). Interestingly, pathway analysis identified the GO terms and KEGG pathways ‘Lysosome,’ ‘Lysosomal Lumen,’ and ‘Lytic Vacuole’ among the top significantly upregulated pathways on the membrane of senescent human fetal lung fibroblasts (IMR-90s), a melanoma cell line (SK-MEL-103), B16-F10 mouse melanoma cells, as well as mouse embryonic fibroblasts (MEFs) (**Figure 1b, c**) where damage-induced senescence (DIS) was induced using chemotherapy drugs (variously doxorubicin, palbociclib or nutlin). This confirms the fundamental role of lysosomal dysfunction in senescent cells, which has also been reported elsewhere (Curnock et al., 2023; Rovira et al., 2022).

**Figure 1.**
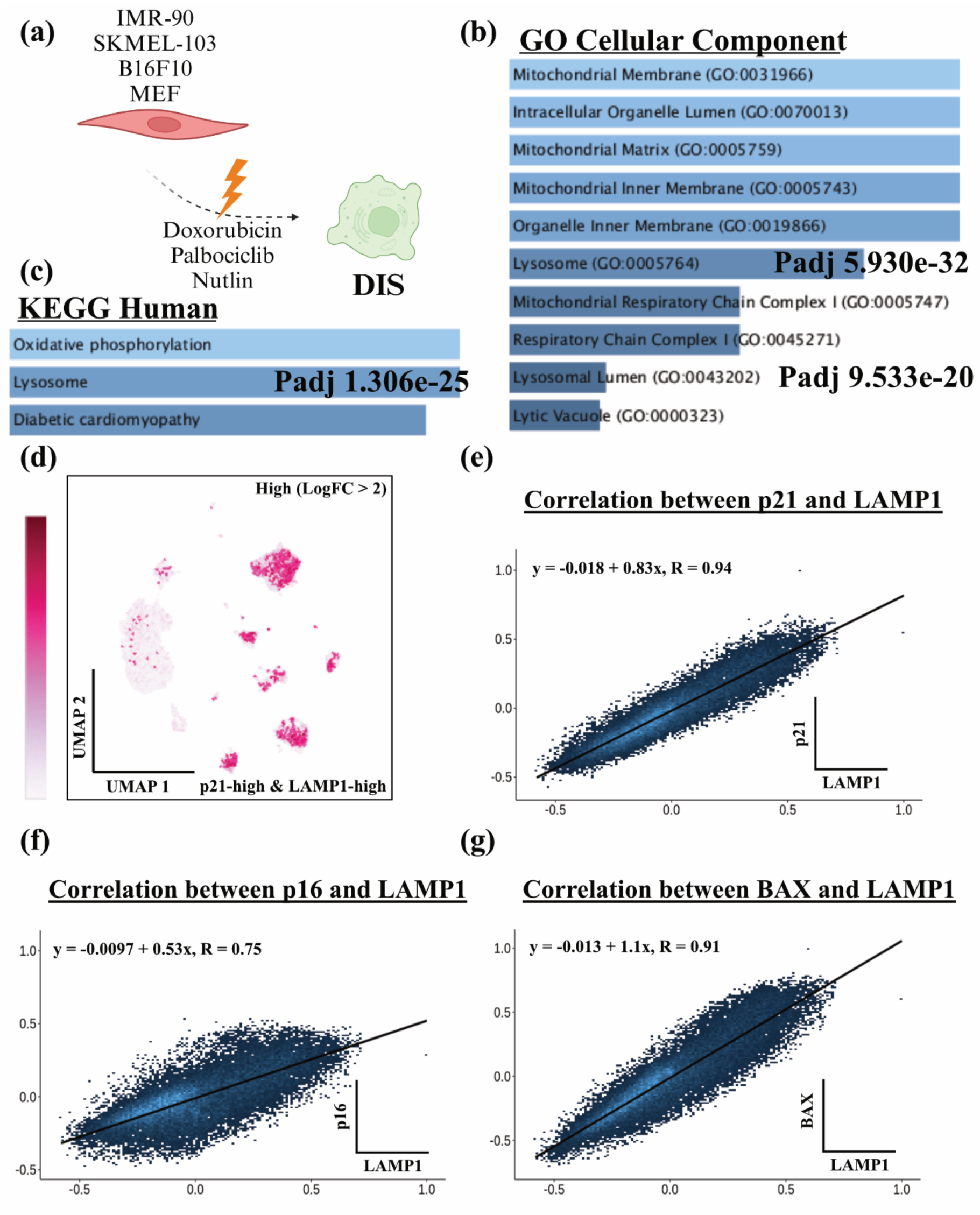
Computational analysis of lysosomal-associated proteins in senescent cells and old tissue. (a) Schematic of the proteomic screen of the plasma membrane of senescent cells. IMR-90, SKMEL-103, B16F10 and MEF cells lines were treated with doxorubicin, palbociclib or nutlin to induce senescence and collected for proteomic analysis (Marin et al., 2023). Created with <u>BioRender.com</u>. (b,c) Pathway enrichment analysis of proteins that were significantly upregulated on the surface of SEN in at least three conditions (Enrichr). (d) Distinct human muscle cells co-expressing high (LogFC > 2) LAMP1 and CDKN1A (p21). (e-g) Correlation between LAMP1 expression and senescence-associated markers (e) p21, (f) p16, and (g) BAX, BCL2 associated X, apoptosis regulator, in healthy tissue (Correlation AnalyzeR).

LAMP1 is essential for lysosomal biogenesis (Rohrer et al., 1996). Analysis of the human muscle aging atlas that consists of almost 300,000 cells isolated from the muscles of young (15 to 46 years old) and old (74 to 99 years old) adults revealed that LAMP1 expression was elevated in multiple cell types in aged cells compared to young (**Supplementary Figure 1a, b**) (Lai et al., 2024). We further observed a strong correlation of LAMP1 with multiple cell populations with high CDKN1A/p21 expression, a critical mediator of cell cycle arrest in SEN (**Figure 1d**). SEN are known to be heterogeneous (Admasu et al., 2021; Hernandez-Segura et al., 2017), so we tested whether this correlation holds true across datasets. We interrogated public data from healthy and cancer tissues and observed a strong correlation of LAMP1 expression with other established senescence genes (Miller & Bishop, 2021). Not only did LAMP1 have a strong expression correlation with p21 (p21 - healthy tissue, R=0.94) (**Figure 1e**), but its expression was also positively correlated with p16 (**Figure 1f**) and BAX (BCL2 associated X, an apoptosis regulator) (**Figure 1g**), which are putative makers of senescence. Moreover, LAMP1 expression strongly correlated with p21 in cancer tissue (R=0.95) and with the surface senescence maker PLAUR (uPAR), which has also been reported as a SEN surface marker and target for senolytic immunotherapy (Amor et al., 2020) (**Supplementary Figure 1c, d**).

### LAMP1 is upregulated on the cell membrane of senescent cells

To confirm the *in silico* transcriptomics and proteomics data, we used a well-characterized doxorubicin model of senescence induction in human fetal lung fibroblasts IMR-90. After 24 hours of treatment with doxorubicin (300 nM), we observed a significant increase in the percentage of SA-β-Gal^+^ IMR-90 fibroblasts (SEN), compared to untreated non-senescent cells (NS) (**Figure 2a, b**). Immunofluorescence confirmed the increased expression of total LAMP1 in permeabilized SEN with negligible detection in NS controls (**Figure 2c**).

**Figure 2.**
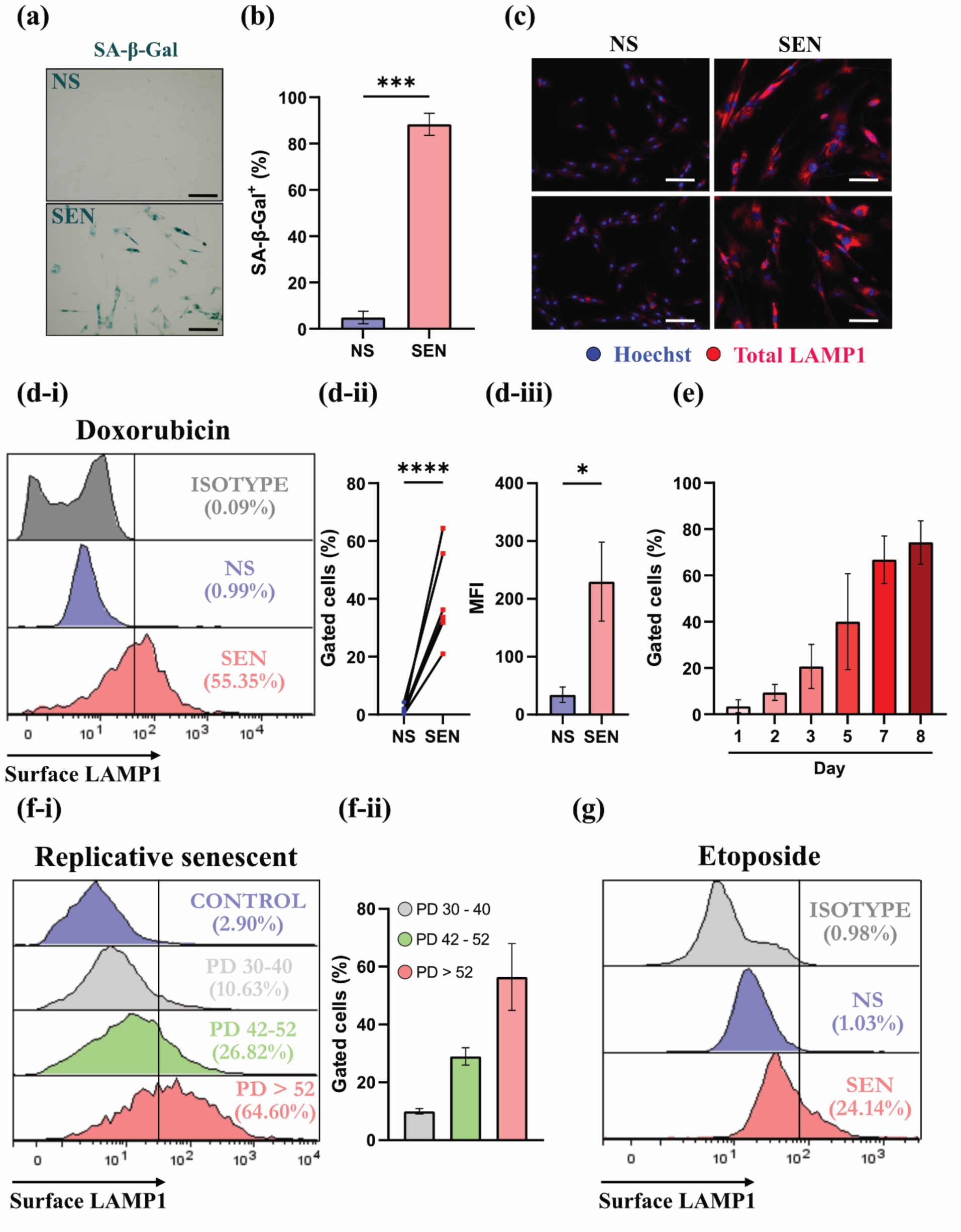
LAMP1 is upregulated on the cellular membrane of senescent cells. (a) SA-β-Gal staining in IMR-90 fibroblasts 9 days after treatment with doxorubicin. Scale bar = 150 μm. (b) Percentage of cells staining positive for SA-β-Gal; n = 3, unpaired t-test; data represented as mean ± S.E.M. (c) Representative images of total LAMP1 staining in SEN doxorubicin-treated fibroblasts and NS controls, n = 3. Red, LAMP1. Blue, Hoechst. Scale bar = 150 μm. (d-i) LAMP1 expression on the surface of SEN and NS cells measured by flow cytometry. (d-ii) LAMP1^+^ percentage of analyzed cells; n ≥ 3, unpaired t-test. (d-iii) Mean Fluorescence Intensity (MFI) of doxorubicin-treated IMR-90 fibroblasts stained for LAMP1, n ≥ 3, unpaired t-test; data represented as mean ± S.E.M. (e) LAMP1^+^ cell proportion in senescent IMR-90 cells at different time points after doxorubicin treatment. n = 2, unpaired t-test; data represented as mean ± SD. (f-i) LAMP1 cell surface staining on IMR-90 cells of increasing PD numbers; n = 2, data represented as mean ± SD. (f-ii) Quantification of gated cells expressing LAMP1 on their cell surface with progressively higher population doubling number. n = 2; data represented as mean ± SD. (g) Representative image (n = 3 biological replicates) of etoposide-treated fibroblasts stained with LAMP1 on the surface of SEN and NS cells. * p ≤ 0.05; ** p ≤ 0.01; *** p ≤ 0.001.

Furthermore, flow cytometric analysis of unfixed and non-permeabilized SEN and NS IMR-90 cells revealed that only about 1% of NS cells express LAMP1 on the surface. In comparison, 20 to 60% of SEN cells express stable surface LAMP1 (**Figure 2d-i, d-ii**). Additionally, SEN showed a five-fold increase in mean fluorescence intensity (MFI) for LAMP1 staining compared to NS control cells (**Figure 2d-iii**). These observations are in line with the *in silico* data showing increased expression of LAMP1 in senescent cells and cells from older individuals.

Previously, Rovira and colleagues reported a gradual increase in LAMP1 gene expression in senescent SK-MEL-103 melanoma cells (Rovira et al., 2022). To determine the kinetics of LAMP1 expression on the surface of SEN, cells treated with doxorubicin for varying number of days were analyzed with flow cytometry. We observed a steady increase in the percent of SEN with surface LAMP1, reaching over 60% by days 7 and 8 after senescence-causing insult (**Figure 2e**).

As the phenotype of SEN may vary depending on the type of initiating damage (Admasu et al., 2021; Hernandez-Segura et al., 2017), we confirmed the surface expression of LAMP1 in other modes of senescence induction. It is well established that continuous passaging of cells results in replicative senescence (Hayflick & Moorhead, 1961). Our results show a progressive increase in surface LAMP1 expression from 10% in cells with PD (population doubling) 30-40, to roughly one quarter in PD 42-52 cells, culminating with more than 60% of replicative senescent cells (PD>52) exhibiting surface LAMP1, versus negligible levels in controls (**Figure 2f-i, f-ii**). Similarly, etoposide is a well-known chemotherapy drug routinely used to induce senescence in cultured cells (Georget et al., 2023). 20% of etoposide-treated IMR-90 cells expressed LAMP1 on their surface, compared to 1% of NS control cells (**Figure 2g, Supplementary Figure 2a, b**).

Given that LAMP1 gene is conserved in mice (mLamp1, or Lamp1), we used doxorubicin to induce senescence in mouse embryonic fibroblasts (MEFs) and NIH/3T3 fibroblasts to confirm our previous observations. MEFs were treated with doxorubicin (150 nM) for 24 hours and senescence was confirmed qualitatively by SA-β-Gal (**Figure 3a**). Flow cytometric analysis of unfixed and non-permeabilized MEFs showed a 10-fold significant increase in the percentage of SEN MEFs expressing surface Lamp1 compared to NS controls (**Figure 3b-i, b-ii**). The analysis of the MFI in MEFs also confirmed a more than 2-fold increase in the Lamp1 expression in the SEN compared to NS cells (**Figure 3b-iii**). Similarly, senescent NIH/3T3 cells increased the expression of Lamp1 on their surface, with 7% Lamp1^+^ cells in the NS population versus 56% in the SEN population (**Supplementary Figure 3a)**. This suggests that cell-surface human and mouse LAMP1 expression is absent or low in NS cells, but it is markedly higher in SEN.

**Figure 3.**
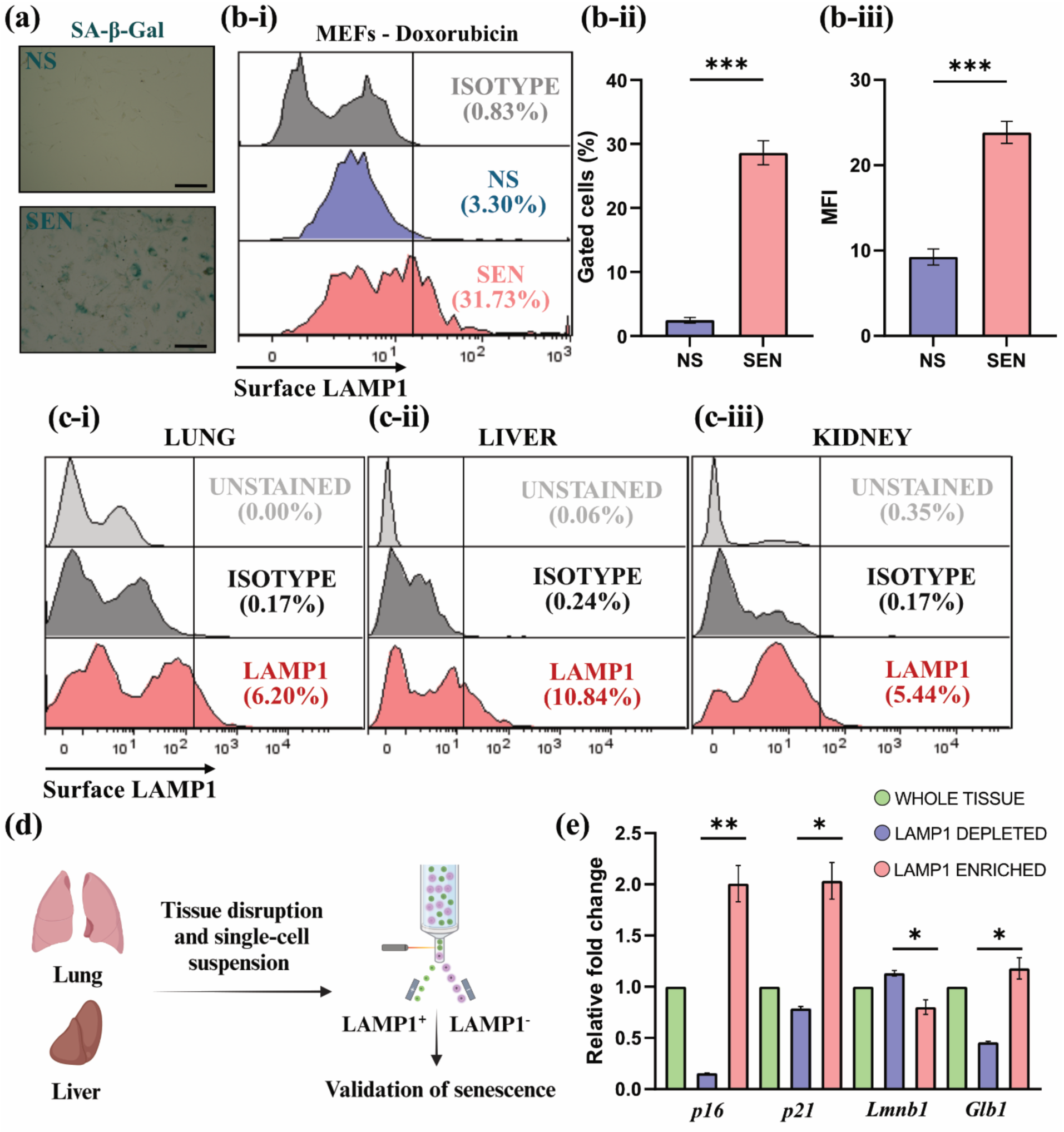
Lamp1^+^ cells express high levels of prototypical senescence markers in mouse tissues. (a) SA-β-Gal staining in MEFs 9 days after treatment with doxorubicin. Scale bar = 150 μm. (b-i) Representative image of Lamp1 expression on the surface of SEN (red) and NS cells (blue) compared to isotype control (grey). Data is representative of n ≥ 3 biological replicates. (b-ii) Quantification of gated cells expressing Lamp1 on their cell surface. Data is representative of n ≥ 3 biological replicates, unpaired t-test; data represented as mean ± S.E.M. (b-iii) MFI of SEN and NS cells stained with Lamp1. Data is representative of n ≥ 3 biological replicates, unpaired t-test; data represented as mean ± S.E.M. (c) Lamp1^+^ population in the lungs (c-i), liver (c-ii), and kidneys (c-iii) of middle-aged mice (39-63 weeks old). Data is representative of n ≥ 2 biological replicates. (d) Schematic of the experimental approach. Single-cells were isolated from mice tissue and sorted based on their Lamp1 status. RNA was isolated and RT-qPCR was used to validate prototypical markers of senescence. Created with <u>BioRender.com</u>. (e) Gene expression of p16, p21, Glb1, and Lmnb1 of Lamp1^-^ and Lamp1^+^ cells. Unsorted single cells were used as controls (green). Data is representative of n ≥ 3 biological replicates, unpaired t-test; data represented as mean ± S.E.M. *Gapdh* and *Actb* were used as housekeeping controls. * p ≤ 0.05; ** p ≤ 0.01; *** p ≤ 0.001.

### Lamp1^+^ cells express high levels of prototypical senescence markers in multiple mouse tissues

The age-dependent increase in senescence burden is well documented (Dimri et al., 1995; Tuttle et al., 2020; Wang et al., 2009). LAMP1 is a widely-expressed lysosomal protein (Eskelinen, 2006). We therefore wondered if the elevated cell surface LAMP1 in SEN that we observed in culture also occurs *in vivo*. We set out to determine cell-surface expression of Lamp1 in the lung, liver, kidneys, and spleen of middle-aged mice (39-63 week-old animals).

Flow cytometry analysis of lung-derived cells reveals only about 6% of cells expressing Lamp1 on their surface (**Figure 3c-i**). Analysis of the liver (**Figure 3c-ii**) and the spleen (**Supplemental Figure 3b**) reveals around 10% of cells expressing cell-surface Lamp1, and only 5% of kidney cells do so (**Figure 3c-iii**). Overall, our results suggest that surface Lamp1 can be detected in multiple mouse organs (**Table 1**), and is expressed in a similar proportion of the cells (1.5 – 10%) as the senescence markers previously reported in the tissues of middle-aged, old, or challenged mice (Wang et al., 2024; Wang et al., 2021; Wang et al., 2009).

**Table 1.** Percentage of Lamp1^+^ cells observed in different mouse organs.

| Organ | Surface Lamp1<sup>+</sup> (%) |
| --- | --- |
| Lung | 3 – 8 |
| Liver | 9 – 11 |
| Kidney | 3 – 6 |
| Spleen | 3 – 11 |
| Bleomycin-treated fibrotic lung | 9 – 24 |

To determine whether the cells positive for surface Lamp1 are indeed senescent, we isolated both Lamp1-enriched (Lamp1^+^) and Lamp1-depleted cell (Lamp1^-^) populations from the liver and lungs of middle-aged mice (**Figure 3d**). We confirmed high purity in the sorted samples (**Supplementary Figure 3c, d**). Consistent with our hypothesis, the quantitative real-time PCR analysis of samples from the liver revealed that cell-surface Lamp1^+^ cells express the prototypical makers of senescence p16 and p21 at higher levels than Lamp1^-^ cells (**Figure 3e**). Notably, p16 expression in the Lamp1 depleted population was even lower than on unsorted cells. Presumably, the whole tissue (unsorted population) contains Lamp1^+^ and Lamp1^-^ cells (**Figure 3e**), leading to the intermediate value. Additionally, Lamp1^+^ cells exhibited features of compromised nuclear integrity, as represented by Lmnb1 downregulation compared to Lamp1^-^ (**Figure 3e**). Finally, Glb1 — the gene encoding lysosomal β-D-galactosidase and responsible for SA-β-Gal activity —is upregulated in cell-surface Lamp1^+^ cells. And in line with our p16 findings, we observed a slight decrease in the expression of Glb1 in Lamp1^-^ compared to whole tissue or Lamp1^+^ cells (**Figure 3e**).

Lamp1^+^ cells from the lung had significantly increased expression of p16 but not p21 compared to Lamp1^-^ cells (**Supplementary Figure 3e**). In line with previous results, lung Lamp1^+^ cells had lower Lmnb1 expression than Lamp1^-^ cells (**Supplementary Figure 3e**). Overall, our results show that, at least in the liver and lung, Lamp1-enriched cells express multiple senescence markers.

### Lamp1^+^ cells accumulate in mice with fibrosis

Our data so far show that not only do cells undergoing senescence in cell culture express LAMP1 on their surface, but that cell surface Lamp1 can be used to enrich cells expressing senescence-associated genes from various tissues. We next investigated whether this strategy could track senescence burden in pathological states in which SEN are implicated. Idiopathic Pulmonary Fibrosis (IPF) is a lung disease characterized by progressive and irreversible scarring (fibrosis) of the lung parenchyma (Juarez et al., 2015). The etiology of IPF remains elusive, but its prevalence is strongly associated with advancing age, particularly in those over 50, with a higher incidence in the 6th and 7th decades of life (Selman et al., 2016). Notably, studies demonstrated increased expression of several senescence markers in various types of cells within the lung of IPF experimental models and patients (Parimon et al., 2021).

It is well-established that the genotoxic agent bleomycin (BLM) induces pulmonary fibrosis in humans (Mohammed et al., 2024) and in the mouse lung (Schafer et al., 2017). In this study we explored whether cell-surface LAMP1 may serve as a biomarker of SEN in mice after IPF induction. As illustrated in **Figure 4a**, bleomycin or saline was administered to mice via the oropharyngeal route. Following bleomycin instillation, the weight of the treated mice decreased, suggesting successful IPF induction (**Figure 4b**). Previously, extensive fibrotic lesions have been reported in the lungs of mice 20-28 days post-bleomycin challenge (Schafer et al., 2017). The picrosirius red stain is one of the most commonly-used histochemical techniques to stain collagen and detect changes in matrix organization characteristic of fibrosis (Vogel et al., 2015). Our data confirmed significantly increased collagen deposition in the lungs of bleomycin-treated mice compared to sham-treated mice (**Figure 4c-i, c-ii**). An increase in TGF-β primarily drives collagen deposition in the IPF lungs (Gimenez et al., 2017). Real-time PCR analysis of the lung tissue confirmed a three-fold increase in the expression of *Tgf-β* in bleomycin-treated compared to sham-treated mice (**Figure 4d**). Additionally, real-time PCR analysis also confirmed an increase in the expression of p21 in lung tissue from bleomycin-treated mice compared to that of sham-treated mice (**Supplementary Figure 4a)**.

**Figure 4.**
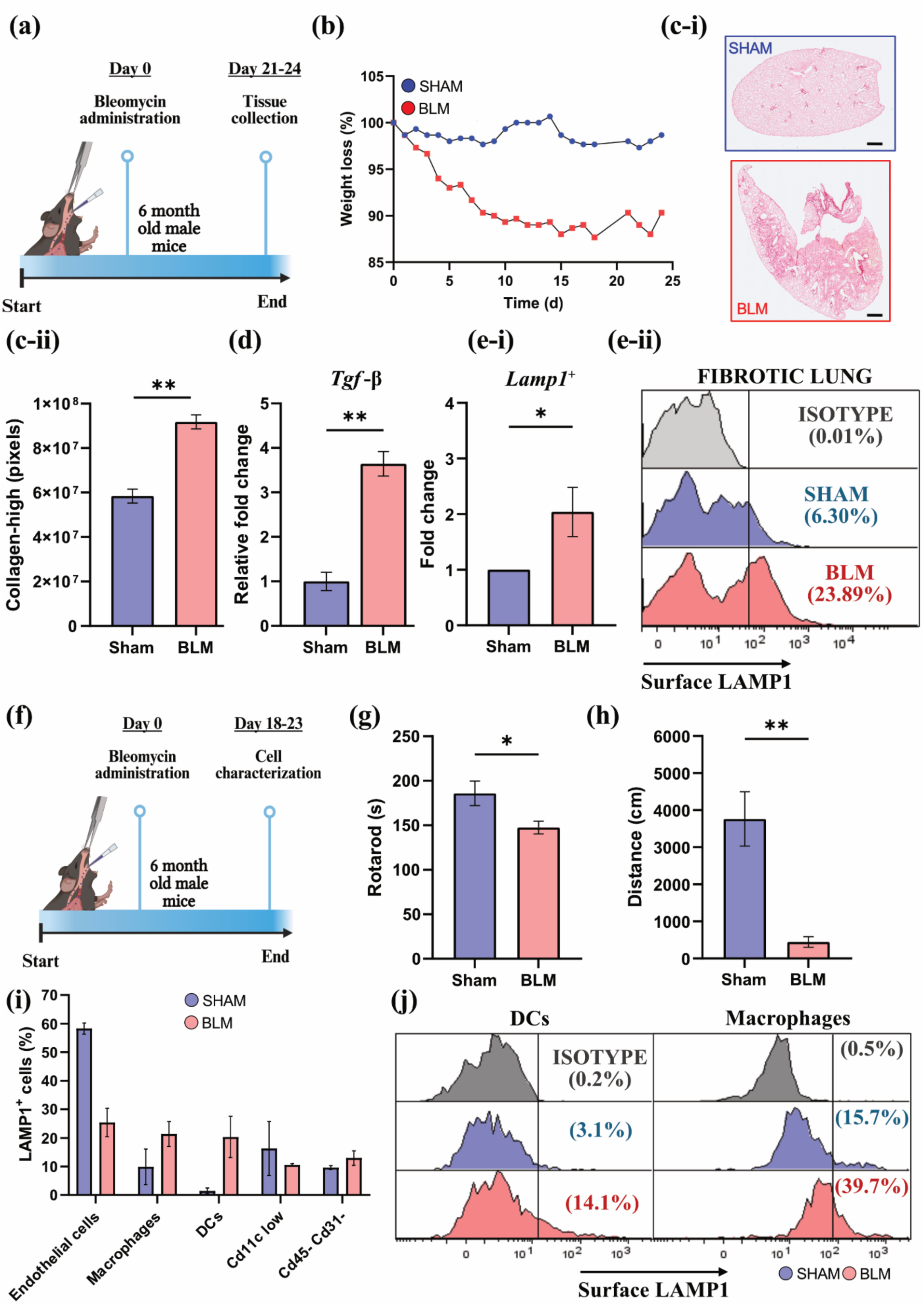
Mice with bleomycin-induced pulmonary fibrosis have a higher percentage and diversity of Lamp1^+^ cells. (a) Schematic of the experimental design for induction of fibrosis and senescence in mice. Bleomycin sulfate was delivered on day 0 using the oropharyngeal route of administration. After 24 days, the lungs were collected. Created with <u>BioRender.com</u>. (b) Bleomycin-induced weight loss. Weight is normalized to the day of bleomycin instillation (day 0), n = 6 (n = 3 saline-treated mice; n = 3 bleomycin-treated mice). (c-i) Representative lung sections stained with Picrosirius red. Scale bar = 500 μm. (c-ii) Quantification of collagen-high areas (pixels) in controls and bleomycin-treated mice (n = 3 saline-treated mice; n = 3 bleomycin-treated mice); data represented as mean ± S.E.M, unpaired t-test. (d) Gene expression of *Tgf-β*. *Hprt* was used as housekeeping control, data represented as mean ± S.E.M, unpaired t-test. (e-i) Fold change increase of Lamp1^+^ cells in the lungs of bleomycin-treated mice compared to saline controls. Data represented as mean ± S.E.M, unpaired t-test. (e-ii) Representative flow cytometry histogram of Lamp1 expression in saline-treated controls and bleomycin-treated fibrotic lungs. Data is representative of n ≥ 3 biological replicates. (f) Schematic of the experimental design for induction of fibrosis and senescence in mice. Bleomycin sulfate was delivered on day 0 using the oropharyngeal route of administration. After 18-23 days, the lungs were collected to characterize cells expressing Lamp1. Mouse behavior was collected on day 17 (n=7; n = 3 saline-treated mice; n = 4 bleomycin-treated mice). Created with <u>BioRender.com</u>. (g) Time spent on a rotarod (seconds) (n=7; n = 3 saline-treated mice; n = 4 bleomycin-treated mice); data represented as mean ± S.E.M, unpaired t-test. (h) Spontaneous movement was measured for 10 minutes during open field (n=7; n = 3 saline-treated mice; n = 4 bleomycin-treated mice); data represented as mean ± S.E.M, unpaired t-test. (i) Characterization of cells expressing surface Lamp1 in the bleomycin-treated fibrotic pooled lungs and the saline-treated mice pooled lungs. Data is representative of n = 3 saline-treated mice, and n = 4 bleomycin-treated mice. (j) Representative flow cytometry histogram of Lamp1 expression in saline-treated controls and bleomycin-treated fibrotic lungs. Live, Cd45^+^, Cd11c^+^, and SiglecF^-^ cells were considered dendritic cells (DCs) (left panel). Live, Cd45^+^, Cd11c^+^, and SiglecF^+^ cells were considered macrophages (right panel). * p ≤ 0.05; ** p ≤ 0.01.

As we previously found that cell surface Lamp1 expression is associated with a senescent phenotype, we next set out to analyze single cells prepared from the lungs of sham- and bleomycin-treated mice. Viable cells from whole-lung single-cell suspensions were analyzed for Lamp1 surface expression (**Supplementary Figure 4c**). Results show a 1.5- to 3-fold increase in the number of Lamp1^+^ cells after the bleomycin challenge relative to sham treatment (**Figure 4e-i, e-ii**). All bleomycin-treated mice showed an increase in Lamp1^+^ cells, but there was mouse-to-mouse variability in the degree of increase (**Figure 4e-i**). Our data points to a greater number of Lamp1^+^ cells in mice with increased senescence burden due to bleomycin treatment.

Previous reports found an increase in the senescence phenotype in different cell types collected from IPF lungs, including endothelial cells (ECs), epithelial cells, AT2 cells, basal cells, fibroblasts, mesenchymal progenitor cells, and innate and adaptive immune cells (Parimon et al., 2021; Schafer et al., 2017). We wondered if an increase in Lamp1 on the cell surface also occurs in multiple cell types in the fibrotic mouse lung. To this end, we repeated the bleomycin installation to induce lung fibrosis (**Figure 4f**). Running time on a rotarod and spontaneous activity were recorded as a measure of general condition deterioration after the treatment, indicative of fibrosis development. Results from the rotarod test reveal a significant decline in the time mice stayed on the rotating rod following bleomycin instillation (**Figure 4g**). The open field test evaluates general activity levels, gross locomotor activity, and exploration habits in mice. Animals were placed into an open arena and allowed to freely explore while data was gathered using an electronic tracking system (Noldus Ethovision). We confirmed a decline in spontaneous activity and the total distance moved in the bleomycin-treated mice compared to sham control mice (**Figure 4h**). To determine which lung cell types express Lamp1 on the surface, we designed a multicolor flow cytometry panel that identifies several cell lineages. Our results show that the majority of Lamp1^+^ cells in the lungs of sham-treated mice were ECs (as defined by Cd45^-^, Cd31^+^) (**Figure 4i**). Small populations of Cd45^+^, Cd11c-high, Siglec-high macrophages, as well as Cd11c-low and undefined Cd45^-^ cells also expressed surface Lamp1 in the sham mice. By contrast, we observed multiple distinct cell types expressing surface Lamp1 after bleomycin instillation. Most of these cells were endothelial cells, macrophages, and dendritic cells (defined as Cd45^+^, Cd11c-high, Siglec-low) (**Figure 4i, j**), while only a small percentage of T cells (Cd45^+^, Cd11c-low, Cd3e^+^) in the bleomycin-treated mice expressed surface Lamp1 (**Supplementary 4c**).

To comprehensively understand the phenotype of cell-surface Lamp1 cells we performed bulk RNA-Sequencing (RNA-Seq.) of Lamp1-enriched, Lamp1-depleted, and unsorted cell suspensions from whole lung in sham-treated and BLM-treated mice 21 days after BLM instillation (**Figure 5a**). We observed a strong increase in fibrotic genes in unsorted lung cells from BLM-treated mice compared to sham. (**Figure 5b**). We compared the upregulated genes in Lamp1-enriched cells in sham-treated mice versus in Lamp1-enriched cells in the bleomycin-treated mice. We observed an overlap of 312 genes between both datasets (**Figure 5c**). Lamp1-enriched cells from sham-treated mice had different pathways upregulated compared to Lamp1-enriched cells from BLM-treated (**Figure 5d, e**). Notably, pathways related to pulmonary fibrosis such as TGF-β regulation, extracellular matrix organization, or collagen biosynthesis were preferentially upregulated in Lamp1-enriched cells from the BLM-treated mice (**Figure 5e**). In line with our hypothesis, Lamp1-enriched cells showed a clear senescence signature compared to Lamp1-depleted cells according to the Sen Mayo gene panel both in sham-treated mice and bleomycin-treated mice (Saul et al., 2022) (**Figure 5f, g**).

**Figure 5.**
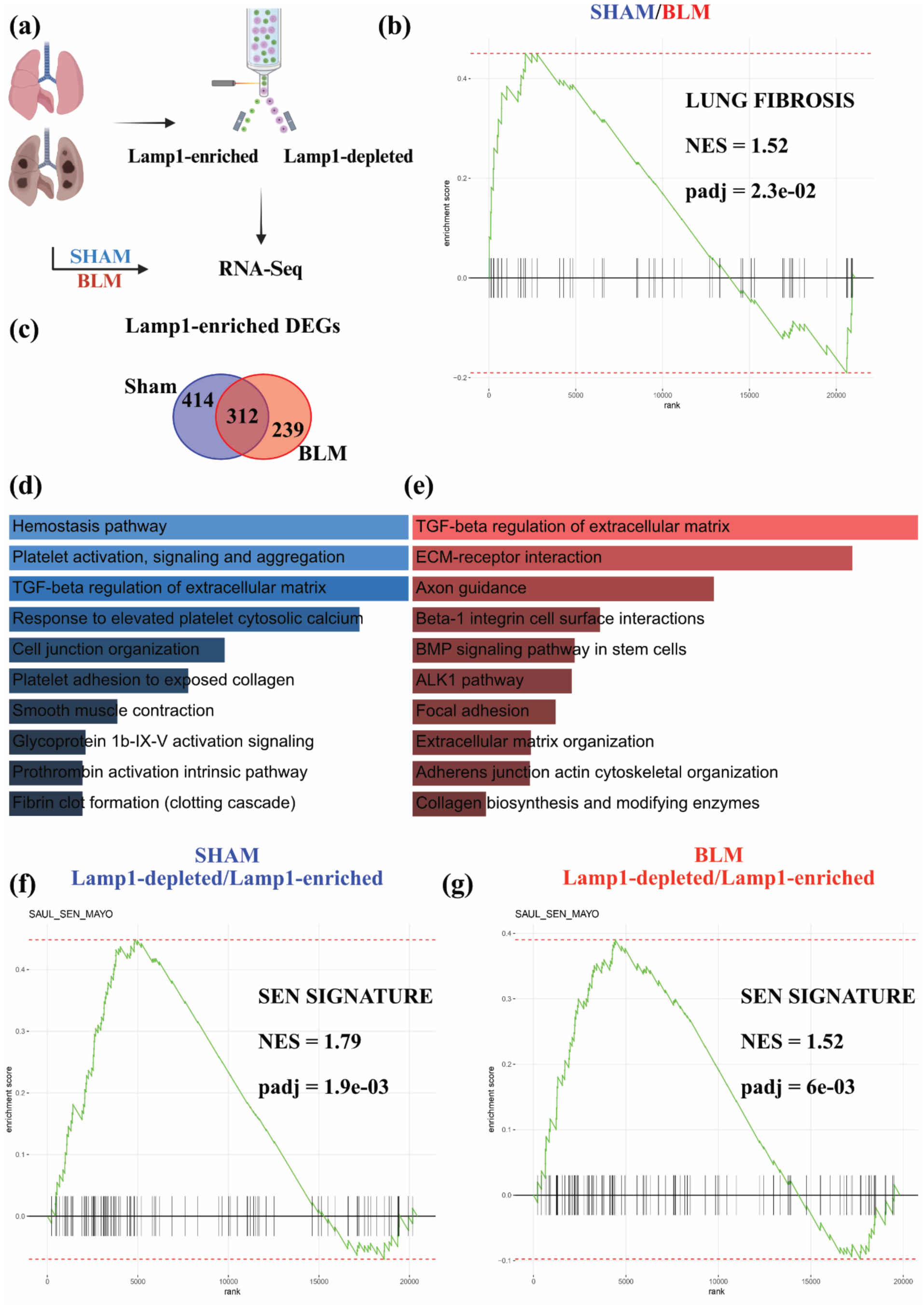
Lung Lamp1-enriched cells have a senescent gene signature. (a) Schematic of the experimental design for induction of fibrosis and senescence in mice. Bleomycin sulfate was delivered on day 0 using the oropharyngeal route of administration. After 21 days, the lungs were harvested and whole tissue single-cells were collected for sequencing. Additionally, cells were sorted based on surface Lamp1 expression and collected for sequencing. Created with <u>BioRender.com</u>. (b) GSEA of sham-treated mice compared to bleomycin-treated mice for the pathway “WP_LUNG_FIBROSIS”. (c) Upregulated genes in the Lamp1-enriched samples in sham- (blue) and BLM-treated (red) mice. Light pink represents genes upregulated in both sham and BLM when comparing Lamp1-enriched versus Lamp1-depleted cells. (d) Upregulated pathways in Lamp1-enriched cells in the sham-treated mice (BioPlanet 2019). All pathways had a padj < 0.05. Pathways on top had lower padj values. (e) Upregulated pathways in Lamp1-enriched cells in the BLM-treated mice (BioPlanet 2019). All pathways had a padj < 0.05. Pathways on top had lower padj values. (f,g) GSEA of sham-treated mice and bleomycin-treated mice for the pathway “SAUL_SEN_MAYO”. Lamp1-depleted cells were compared to Lamp1-enriched cells for (f) sham-treated mice, and (g) BLM-treated mice.

### A LAMP1-targeting antibody-drug conjugate (ADC) targets senescent cells

Finally, we wanted to assess whether SEN cell-surface LAMP1 can be exploited to target and clear SEN. First, to test LAMP1 surface expression in physiological conditions, we performed LAMP1 labeling for flow cytometry at both 4°C and 37°C. We observed that LAMP1 remained stable and selectively elevated on the surface of SEN in both conditions (**Figure 6a**). We then utilized an antibody-drug conjugate (ADC) approach to test if cell-surface LAMP1 can be targeted to selectively deliver a cytotoxic payload to SEN. SEN and NS controls were incubated with either a human anti-LAMP1 antibody or an IgG control antibody. Following incubation, a secondary ADC antibody targeting the primary anti-LAMP1 antibody and conjugated with a cytotoxic duocarmycin payload was added to the wells. To assess cell viability, we used the xCELLigence RTCA-MP (Agilent, USA), which tracks real-time cell viability by measuring the cellular impedance of live cells. Changes in impedance are expressed as a cell index (CI) value, which derives from relative impedance changes corresponding to cellular coverage of the electrode sensors, normalized to baseline impedance values with medium only. Percent cytotoxicity was determined based on the normalized CI value of cells treated with the antibody-ADC sequence relative to control cells untreated with either. Results show a significant decline in the normalized CI in SEN treated with the two-part anti-LAMP1 ADC strategy compared to the cells treated with the IgG control antibody plus ADC. In contrast, this treatment did not affect the viability of the NS control cells (**Figure 6b**). Our results further show an increase in percent cytotoxicity in SEN following treatment with the two-part ADC within 10 hours and a steady increase in the cytotoxicity to around 50% by the 48-hour mark. We did not observe any increase in cytotoxicity in NS cells treated with this approach (**Figure 6b, c, d**). We conclude that, senescent cells can be targeted using an anti-LAMP1 ADC treatment (**Figure 6e**).

**Figure 6.**
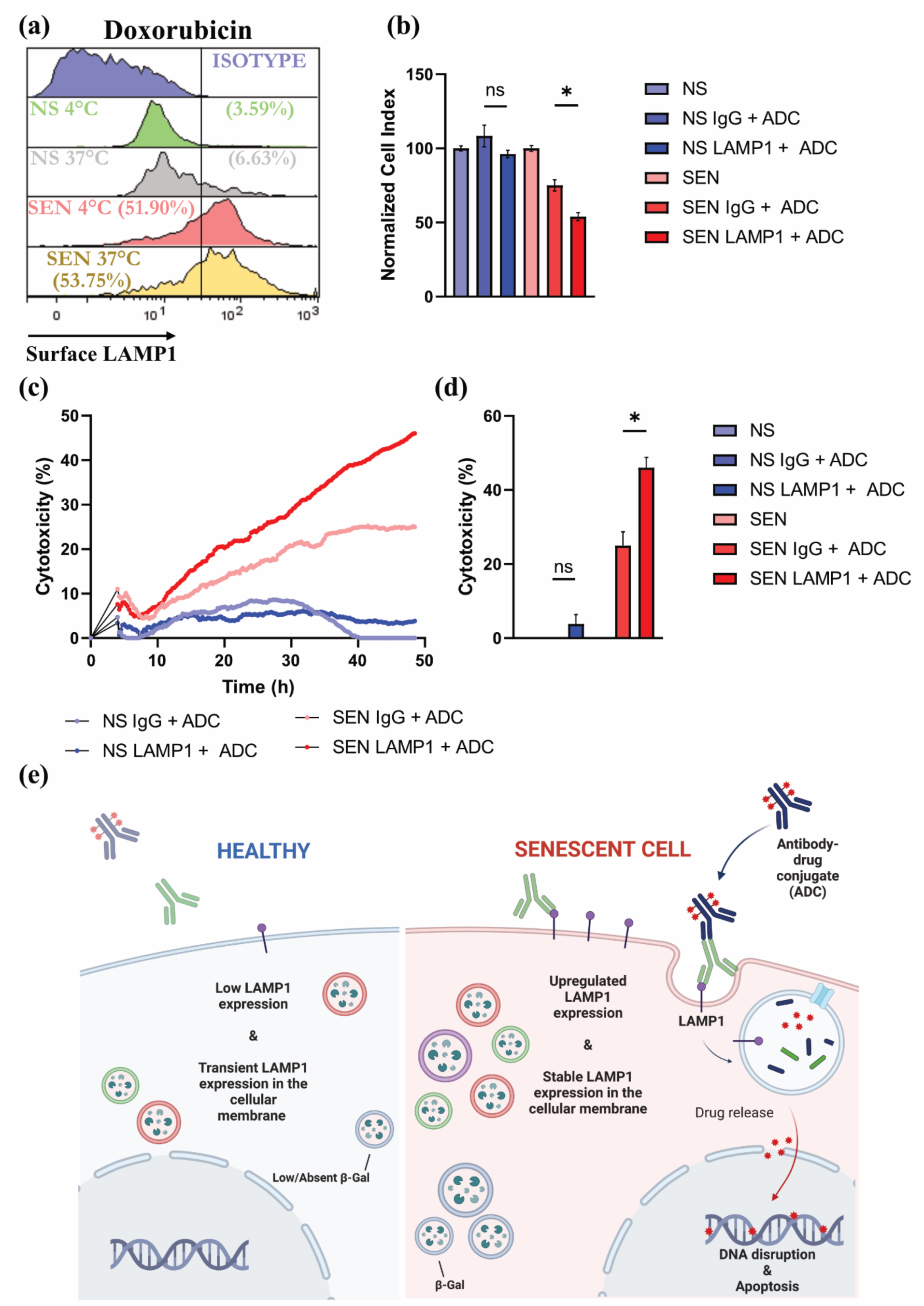
A LAMP1-ADC kills senescent cells. (a) LAMP1 expression on the surface of SEN and NS cells at 4°C and 37°C. Data is representative of n ≥ 3 biological replicates. (b) Normalized cell index in SEN and NS after 48 of labelling with LAMP1 plus ADC, IgG control plus ADC, or nothing. Data is representative of n ≥ 3 biological replicates; mean ± S.E.M, unpaired t-test. (c) Cytotoxicity in SEN and NS after 48 of labelling with LAMP1 plus ADC, IgG control plus ADC, or nothing. Data is representative of n ≥ 3 biological replicates; mean ± S.E.M, unpaired t-test. (d) Real time kinetics of ADC killing NS and SEN cells as measured by the xCelligence RTCA platform every 15 minutes. Untreated NS and SEN were used as controls (0% death). Data is representative of n ≥ 3 biological replicates. (e) Antibody-drug conjugate approach to target senescent cells while sparing healthy non-senescent cells schematic. Created with <u>BioRender.com</u>.

## Discussion

Identifying senescent cells in living tissue based on the activity of SA-β-Gal has been critical for understanding the biology of aging and the role of cellular senescence in physiology (Dimri et al., 1995). Since then, several other markers have been used to identify senescent cells, including increased expression of the cell cycle checkpoint inhibitors p16 and p21, SASP factors such as IL-6, IL-8 and MMP9, and release of the DAMP signaling molecule HMGB1 into the extracellular milieu (Hernandez-Segura et al., 2018). Use of these markers has enabled the discovery of the role of senescent cells in the biological aging process and in diseases of aging, and has identified SEN as targets for treatment of numerous diseases of aging and for degenerative aging writ large (Kirkland & Tchkonia, 2017). However, no known ‘unique’ senescence markers exist, and most existing markers are useful in cell culture models but have limited application in the identification of senescence in tissues or require transgenic models that cannot be translated to human medicine. The lack of specific and broadly applicable markers of senescent cells in tissues and living organisms has led to guidelines called ‘minimum information for cellular senescence experimentation *in vivo*’ (MICSE) that recommends measuring at least two or more makers for greater confidence (Ogrodnik et al., 2024).

Here, we build upon prior findings that lysosomal dysfunction is characteristic of cellular senescence and is often associated with aging and age-related diseases (Curnock et al., 2023; Rovira et al., 2022). Analysis of previously published single-cell RNA Sequencing (scRNA-Seq) data shows a strong correlation of expression of the lysosomal membrane protein LAMP1 with p21. This was further confirmed by studying the correlation of LAMP1 with several senescence phenotypes in healthy and cancer tissue samples, suggesting its involvement with the senescence cell fate. Increased expression of lysosomal proteins during senescence has been previously reported (Curnock et al., 2023; Deng et al., 2024; Kang et al., 2017; Marin et al., 2023; Rovira et al., 2022). In agreement with these studies, we observed an increase in the expression of total LAMP1 in senescent cells by immunofluorescence assay. While no previous studies have focused on LAMP1 as a selective and sensitive marker of senescence, it has been identified among over-represented cell surface proteins in several prior reports (Deng et al., 2024; Marin et al., 2023). A surfaceome mass spectrometry screen of NHLFs, MEFs, primary mouse astrocytes, and HUVECs induced to senesce using several methods (genotoxic stress, oxidative stress, or proteasome-induced stress) revealed that LAMP1 was among the top 27 differentially-expressed protein (DEPs) in the membrane of senescent cells (Deng et al., 2024). An increase in the LAMP1 expression has also been previously reported on the surface of senescent cells from the human melanoma line SK-MEL-103 (Rovira et al., 2022) and in the membrane fraction of etoposide-treated senescent cells (Kang et al., 2017).

In this study, we observed about 10% of cells that were Lamp1^+^ in the liver of 39-63 weeks old mice, which is similar to the 8.4±1.7% of hepatocytes deemed senescent in mice of a similar age via γH2AX foci (Wang et al., 2009) and expectedly lower than the 15% of hepatocytes deemed senescent in 24-month-old mice using TAF (Ogrodnik et al., 2017). The quantitative real-time PCR analysis of Lamp1^+^ cells further confirmed that expression of several other senescence makers (p16, p21, and Glb1) was elevated in these cells compared to Lamp1^-^ cells, with the unsorted mixed population being intermediate between the two.

Our results show that around 6% of cells in the lung of middle-aged mice express Lamp1 on the surface, which not only had significantly higher *p16* but also a lower *Lmnb1* expression compared to the cells without surface Lamp1. Similarly, Wang et al. reported 6.7% SEN based on TAF in the lungs of 12-month-old mice (Wang et al., 2009), and Biran et al. reported about 6-7% SA-β-Gal^+^ cells in the lungs of older (24-month-old) mice (Biran et al., 2017). An increase in senescence burden is a characteristic of several pathologies, including inflammatory lung diseases like IPF and COPD (Schafer et al., 2017). An increase in the expression of SA-β-Gal, p16, or p21 in the lungs of mice treated with bleomycin is also well documented (Biran et al., 2017; Schafer et al., 2017). Consistent with these findings, our data from the bleomycin-treated mouse model of IPF also confirmed a significant increase in the proportion of cells in the lung expressing Lamp1 on the surface, with a high degree of mouse-to-mouse variability (9-24%).

Lamp1 is also known to be transiently expressed on the surface of many cell types, most notably immune cells such as activated NK and T cells (Cohnen et al., 2013). In the bleomycin-treated mice, cells expressing surface Lamp1 were predominantly macrophages, and endothelial cells, with smaller numbers of other types of cells like dendritic cells. In sham-treated mice, most cells that expressed Lamp1 on their surface were endothelial cells. Others have previously shown a senescent phenotype in different types of cells in the IPF lungs (Parimon et al., 2021). In contrast, we did not observe a change in the proportion of fibroblasts or epithelial cells in sham versus bleomycin-treated mice lungs based on surface Lamp1 expression, even though AT2 cells in the lung are often associated with increased senescence burden (Yao et al., 2021). Mesenchymal stem cells (MSCs) express a significant amount of the protein Lamp1 on their cell surface when undergoing differentiation into adipocytes. However, in the lung this is a rare population (Xu et al., 2020). Interestingly, a recent study using genetic tools to trace the functional role of p16-expressing cells reported that senescent macrophages and ECs represent distinct cell populations with different fates and functions during liver fibrosis and repair as removal of p16-expressing macrophages significantly reduced tissue damage, whereas p16-ECs were involved in tissue repair and regeneration (Zhao et al., 2024). In line with this, we observed that most Lamp1^+^ cells were endothelial cells in the sham-treated mice. After bleomycin-challenge, we observed an increase in macrophage-expressing cell-surface Lamp1. This finding invites more investigations into the role of senescence in different cell types in the pathology of IPF. Whether this phenomenon exists in other age-related pathologies must be studied in the future. Also, it presents an interesting possibility to test if selective delivery of senolytic interventions to the type of cell (like senescent macrophages instead of senescent endothelial cells) may further fine-tune the efficacy of senolytic interventions in the resolution of IPF. The RNA-Seq. analysis of the Lamp1-enriched populations in sham and BLM mice lung tissue revealed enrichment of several senescence-related genes in both groups when compared to the SEN_MAYO gene set derived from transcriptomic profiling of senescence markers in Mayo Clinic research datasets further confirming senescence phenotype in the Lamp1-enriched populations.

Recent studies have investigated immune surveillance of senescent cells in aging and disease (Burton & Stolzing, 2018; Pereira et al., 2019; Sagiv et al., 2016; Zhang et al., 2022). These efforts have led to the development of immune-based therapeutic interventions for treating age-related diseases (Arora et al., 2021; Yang et al., 2023). The discovery and validation of several SEN cell-surface markers, including ApoD, DPP4, and PLAUR (uPAR), opens the door to clearing SEN by targeting these antigens using engineered lymphocytes (Amor et al., 2020; Burton & Stolzing, 2018; Kim et al., 2017; Poblocka et al., 2021; Rossi & Abdelmohsen, 2021; Takaya et al., 2023; Yang et al., 2023). uPAR-specific CAR T cells efficiently ablate senescent cells *in vitro* and *in vivo*, extend the survival of mice with lung adenocarcinoma treated with senescence-inducing chemotherapy, and restore tissue homeostasis in mice in which liver fibrosis is induced chemically or by diet (Amor et al., 2020). Moreover, treatment with anti-uPAR CAR T cells ameliorate age-associated metabolic dysfunction, for example, improving glucose tolerance, in aged mice and in mice on a high-fat diet (Amor et al., 2024). However, CAR-T therapeutic interventions are expensive and not free from potential adverse effects (Chohan et al., 2023).

Here, we performed proof of principle experiments to test the therapeutic targeting of senescent cells based on the expression of surface LAMP1 using a two-part ADC. Others have tried similar approaches with other surface makers, including ApoD (Takaya et al., 2023) and DPP4 (Kim et al., 2017). The results from cell culture data were promising and should be tested *in vivo* in the future.

Our results demonstrated that several types of senescent cells express LAMP1 on the surface. However, the heterogeneity of the senescence phenotype requires further investigation. While initially thought to be a uniform phenotype, recent research has revealed significant variability in the senescence phenotype, driven by factors such as cell type of origin, cell cycle, and the nature of the inducing stimulus (Admasu et al., 2021; Hernandez-Segura et al., 2017). Advances in scRNA-Seq and other -omic technologies enable the detailed characterization of SEN heterogeneity at the single-cell level, revealing diverse characteristics and behaviors of senescent cells within a population and providing insights into the underlying molecular mechanisms (Cohn et al., 2023). Future research will allow a better understanding of the diversity of Lamp1^+^ cells in the lung and other organs and shed light on their distinct roles in driving aging and age-related disease.

Another interesting question that remains to be answered is the mechanism by which LAMP1 is presented on the surface of senescent cells. It has been hypothesized that proteins from the lysosome and other phospholipid-rich bilayer organelles could be stably present in the plasma membrane as a mechanism to repair or support the damaged cell surface (Corrotte et al., 2015; Reddy et al., 2001; Sarafian et al., 1998). When the cellular surface is damaged, lysosomes may fuse to the plasma membrane to stabilize the lipid bilayer. This hypothesis is supported by a report from Reddy and colleagues who described a Ca^2+^-regulated exocytosis mechanism that mediates plasma membrane damage repair (Corrotte et al., 2015; Idone et al., 2008).

LAMP1 is typically localized on lysosomal membranes, but it can also be transiently displayed on the cell surface under certain conditions, which is of particular interest in immunology and cancer research due to its role in cell signaling, migration, and immune modulation (Krzewski et al., 2013). For instance, in immune cells, such as T cells and natural killer (NK) cells, LAMP1 is commonly translocated to the cell surface as a marker of degranulation or cellular activation. This happens when cytotoxic granules (lysosome-like organelles) fuse with the plasma membrane, displaying LAMP1 and other lysosomal proteins on the cell surface. LAMP1 can also reach the cell surface via vesicular transport from the trans-Golgi network (TGN) or endosomes. After being synthesized and glycosylated in the endoplasmic reticulum (ER) and Golgi, LAMP1 is directed to lysosomes through clathrin-coated vesicles with mannose-6-phosphate receptors or other sorting receptors. However, a small fraction of LAMP1 can bypass the lysosomes and be trafficked directly to the plasma membrane through secretory vesicles, particularly under conditions that impair normal lysosomal trafficking. While these are examples of transient processes by which LAMP1 may be briefly displayed on the cell surfaces under homeostatic conditions, many tumor cells show a persistent increase in surface expression of LAMP1, which has been linked to their ability to evade immune detection and metastasize. The expression can be upregulated under cellular stress, such as hypoxia or nutrient deprivation, which induces autophagy and lysosomal biogenesis. These conditions increase lysosomal exocytosis and result in greater LAMP1 display on the surface (Agarwal et al., 2015). This aberrant increase in cell-surface LAMP1 expression is more reminiscent of the stable presence of LAMP1 on the cell surface of SEN that we report here and awaits further investigation.

In conclusion, our results show that LAMP1 is upregulated on the surface of SEN compared to NS controls. LAMP1 is upregulated on the membrane in human cells induced to senesce in a variety of ways. Similarly, Lamp1 is upregulated in multiple senescent mouse cell lines. We show that SEN can be sorted from tissues using the expression of Lamp1 on the cell surface. Finally, we confirmed our findings *in vivo* using a fibrosis mouse model where we found greater abundance of Lamp1^+^ cells. In summary, we propose lysosomal-associated membrane proteins, particularly LAMP1, as selective markers of cellular senescence *in vitro* and *in vivo*.

---

**Supplementary Figure 1.**
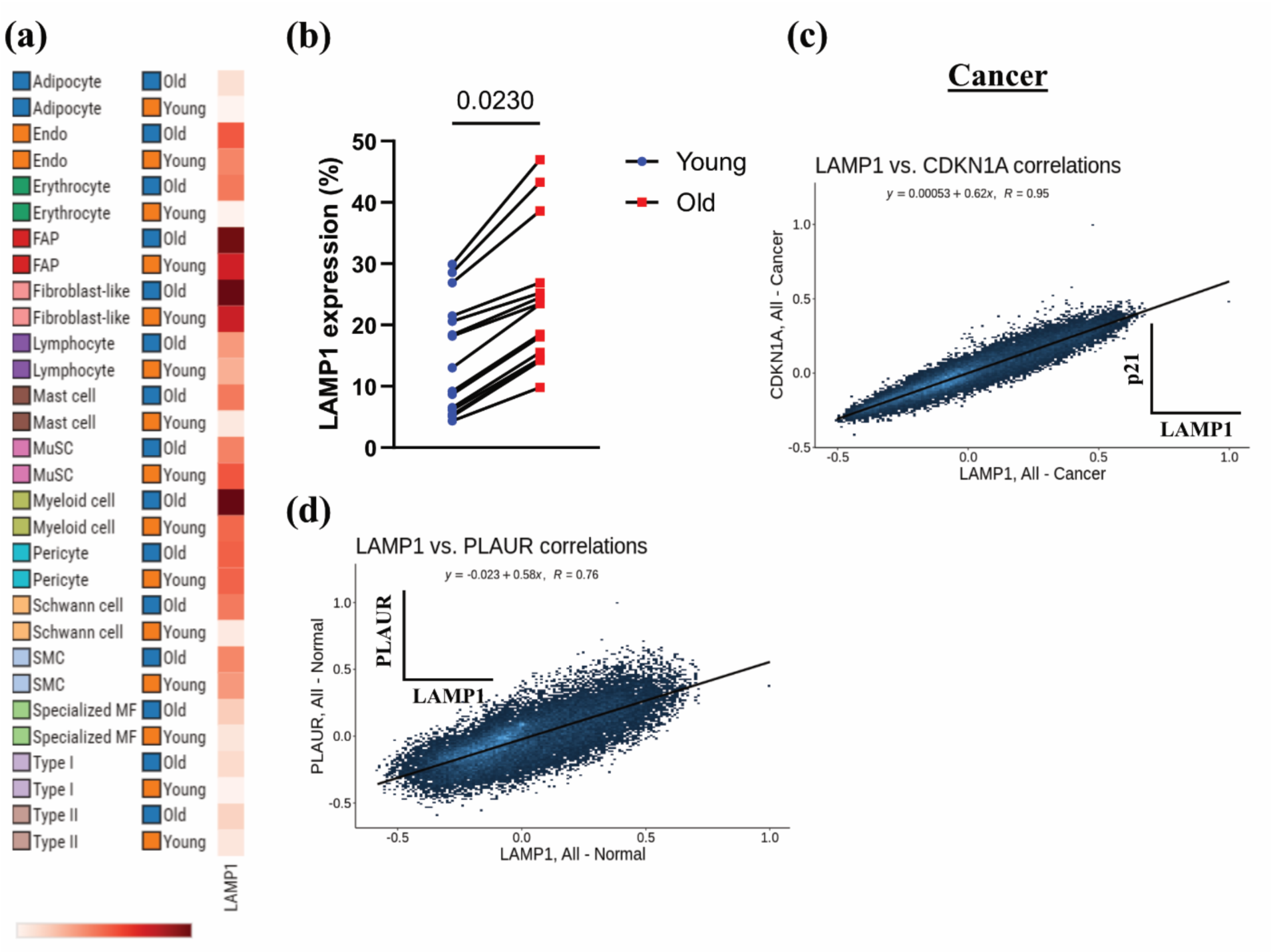
LAMP1 expression increases with age and correlates with senescence-associated genes in healthy and cancer tissues. (a,b) LAMP1 expression in human muscle cells from young and old donors. Data acquired from Muscle Cell Aging Atlas (Lai et al., 2024). (c) LAMP1 expression correlation with CDKN1A (p21) expression in cancer tissue (Correlation AnalyzeR). (d) LAMP1 expression correlation with the senescence surface biomarker uPAR (PLAUR) in healthy tissue (Correlation AnalyzeR).

**Supplementary Figure 2.**
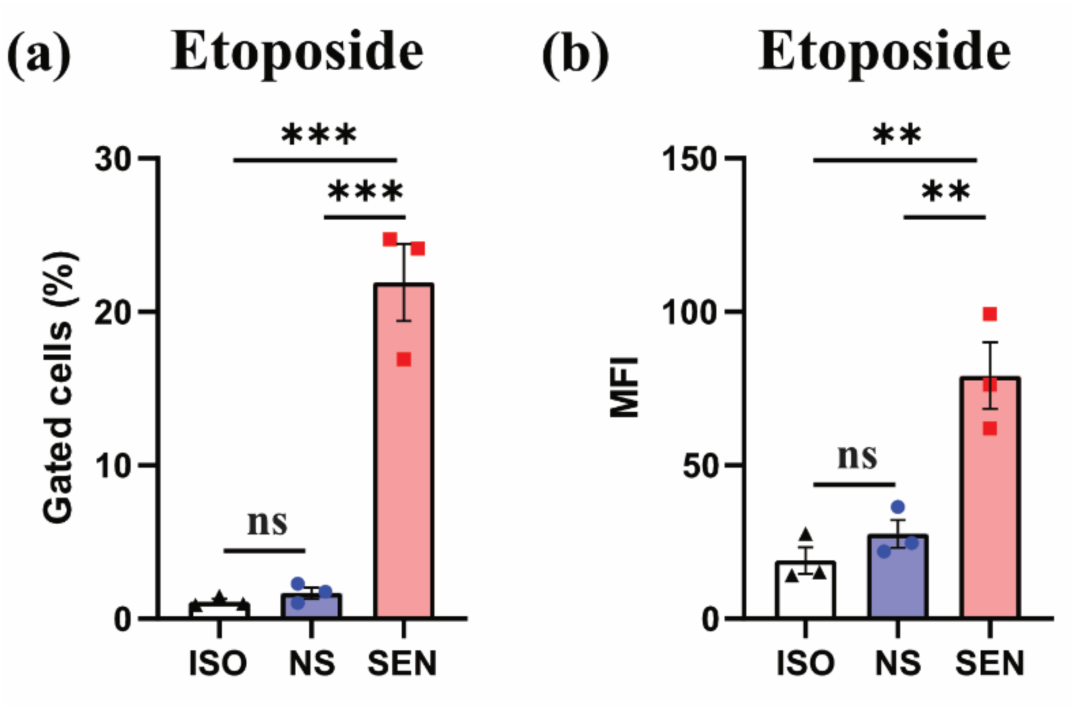
Etoposide-induced senescent cells upregulate LAMP1 on the plasma membrane. (a) Percentage of LAMP1^+^ cells in total cell population following treatment with etoposide, n = 3 biological replicates, unpaired t-test. (b) MFI of etoposide-treated IMR-90 fibroblasts stained with LAMP1, n = 3 biological replicates, unpaired t-test; data represented as mean ± S.E.M. ISO = Isotype control. ns p > 0.05; ** p ≤ 0.01; *** p ≤ 0.001.

**Supplementary Figure 3.**
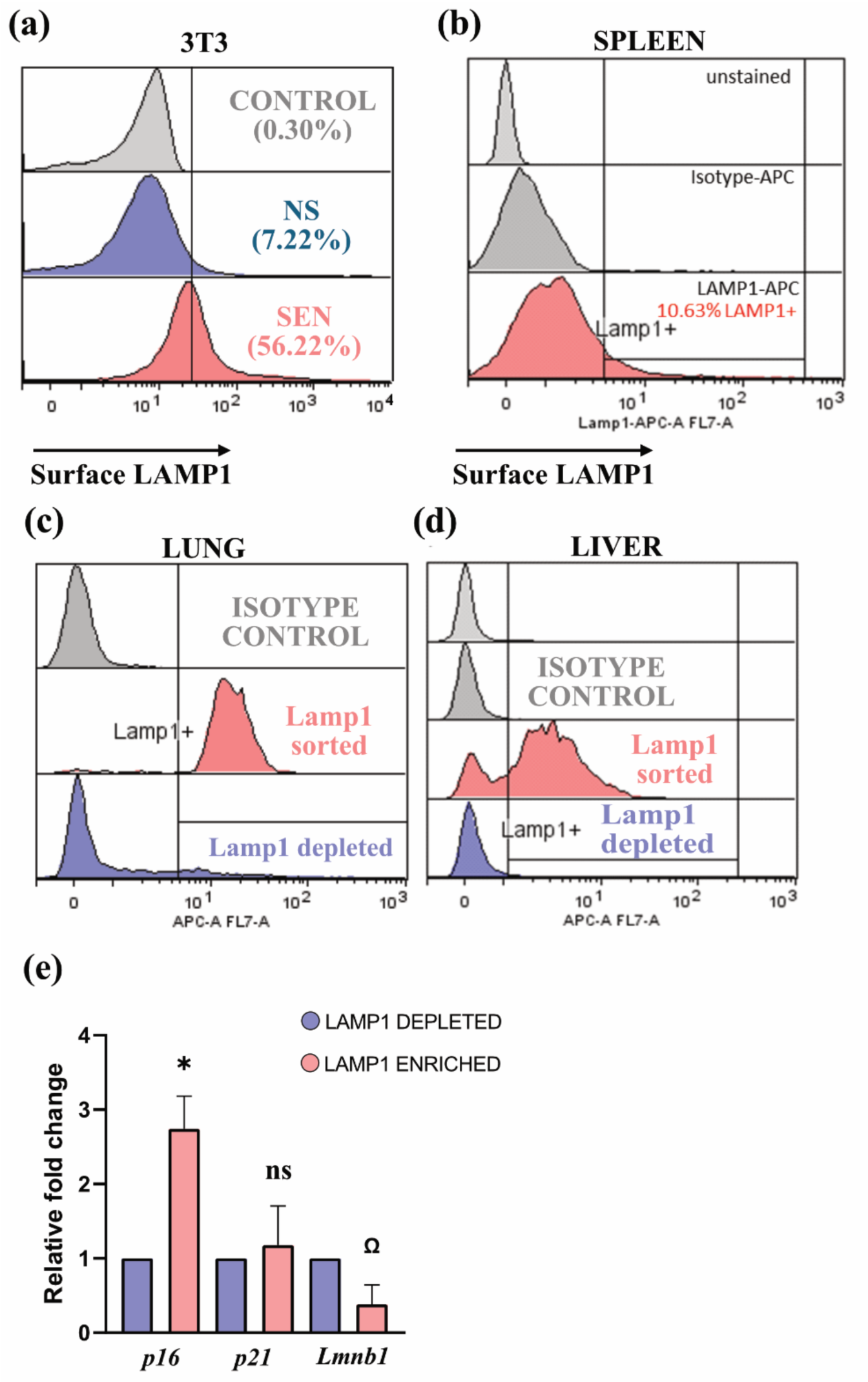
Lamp1 expression in mouse senescent cells and organs. (a) Representative flow cytometry histogram of Lamp1 cell surface expression in mouse 3T3 cells. Light blue, NS controls. Light red, SEN. Data is representative of n = 2 biological replicates. (b) Lamp1^+^ cells in the spleen of middle-aged mice (39-63 weeks old). Data is representative of n = 2 biological replicates. Light grey, unstained. Black, isotype control. Red, Lamp1 expression. (c, d) Cell fractions sorted for LAMP1 have an increased LAMP1 cell surface expression in (c) lungs and (d) liver. Light red, cells sorted based on Lamp1 expression. Data is representative of n = 3 biological replicates. (e) Gene expression of *p16*, *p21*, and *Lmnb1* of Lamp1^-^ and Lamp1^+^ cells. Lamp1-depleted live single cells were used as controls (blue). Data is representative of n ≥ 3 biological replicates, unpaired t-test; data represented as mean ± S.E.M. *Gapdh* used as housekeeping controls. ns p > 0.05; *p<0.05; Ω p<0.08

**Supplementary Figure 4.**
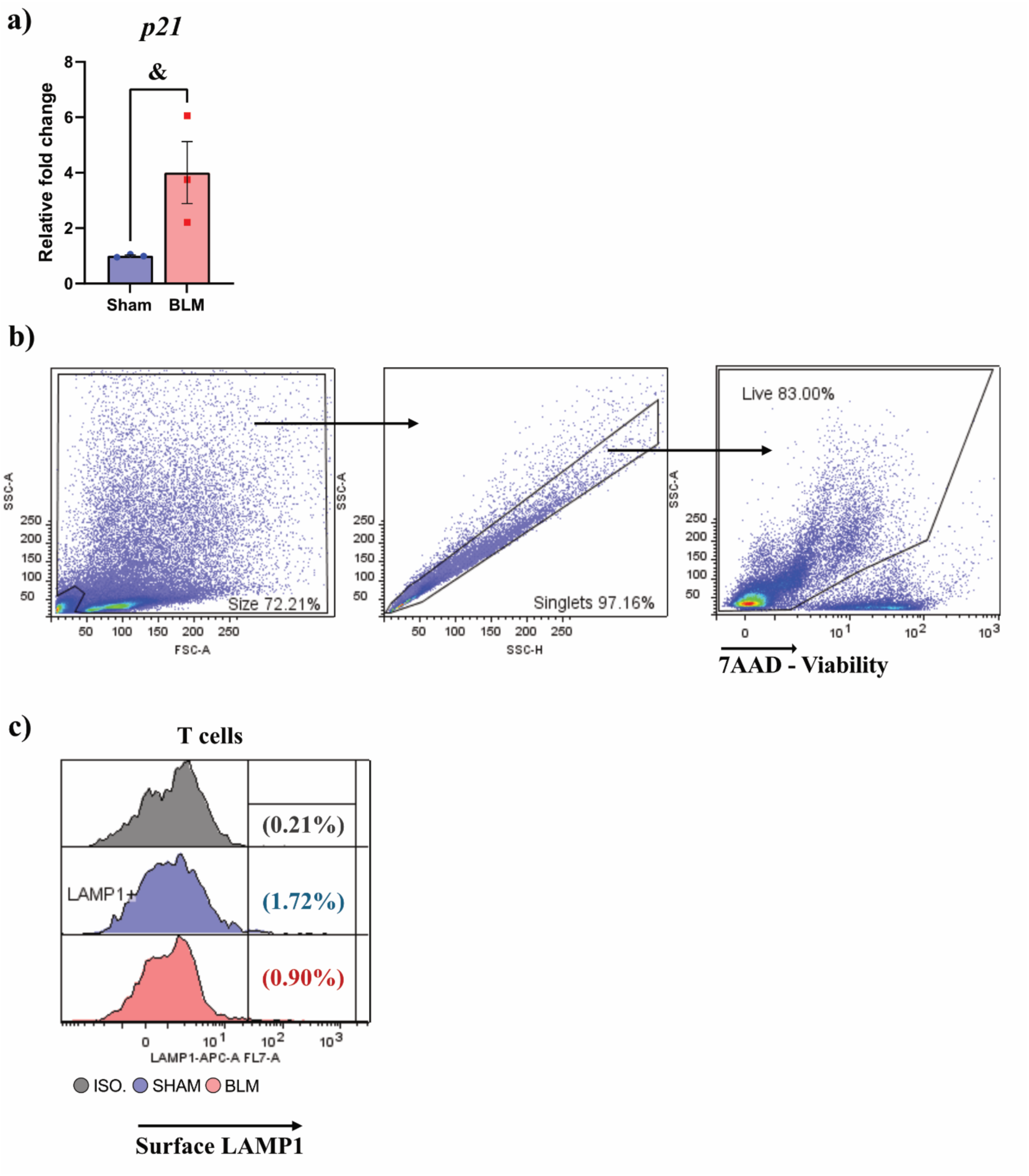
Validation of fibrosis and senescence induction after bleomycin. (a) Gene expression of *p21*. *Hprt* was used as a housekeeping control, data represented as mean ± S.E.M, unpaired t-test. & p<0.06 (b) Gating strategy used to isolate Lamp1^+^ cells. (c) Representative flow cytometry histogram of Lamp1 expression in saline-treated controls and bleomycin-treated fibrotic lungs. Cd45^+^, Cd11c^-^, and Cd3^+^ cells were considered T cells.

## Materials and Methods

### Cell culture and senescence induction

IMR-90 (human lung fibroblasts) and MEFs (mouse embryonic fibroblasts) were cultured in DMEM medium (Corning) supplemented with 10% fetal bovine serum (Sigma) and 1X Penicillin-Streptomycin (Corning) in a humidified atmosphere with 5% CO_2_, 3% O_2_ at 37°C. Cells were routinely checked for mycoplasma.

To induce senescence *in vitro*, IMR-90 cells were treated with 300 nM doxorubicin (Tocris) for 24 hours, then kept in regular culture medium for 9 days before cell collection. Senescence establishment was evaluated ten days after doxorubicin treatment. MEFs received 150 nM of doxorubicin for 24 hours and were considered senescent after nine days post-challenge. To achieve replicative senescence, IMR-90 were continually passaged until exhaustion of replication. Replicative senescence was confirmed by SA-β-Gal assay.

### Senescence-Associated β-Galactosidase

SA-β-Gal staining was performed using a Senescence Detection Kit (Abcam) following manufacturer instructions. Shortly, cells were fixed using a fixative solution, washed, and stained with a staining solution containing 20 mg/mL X-Gal. Cells were incubated for 24 hours, without CO_2_, and imaged.

### Animals

All animal protocols were reviewed and approved by the IACUC. Male C57BL/6J mice used for modeling idiopathic pulmonary fibrosis were purchased from the Jackson Laboratory. All mice were given food and water ad libitum. To induce pulmonary fibrosis, 3 or 3.5 - 4 mg/kg of Bleomycin sulfate (MedChem Express) was administered using the oropharyngeal administration route. Mice were weighed daily and assessed for well-being by a trained specialist. Mice were given additional food and gels to mitigate bleomycin-induced weight loss. Lungs were collected 18-24 days after bleomycin instillation. For validation of fibrosis, the left lobe of the lung was collected for histology, and the right lobes were immediately flash frozen and used for RT-qPCR. For flow cytometry, whole lungs or left and right lobes were dissociated and stained as described below.

### Histology

Lungs were harvested and fixed in 10% neutral buffered formalin for 24 hours before being transferred to ethanol 70% prior to paraffin embedding. 5 μm paraffin-embedded sections were deparaffinized, rehydrated and treated with 0.2% phosphomolybdic acid aqueous solution for 5 minutes. Sections were then stained with Picrosirius red prepared in 0.1% saturated aqueous solution of picric acid for 2 hours at room temperature. The sections were washed with 0.015N hydrochloric acid for 3 minutes, rinsed in 70% ethanol, dehydrated with increasing concentrations of EtOH and xylene. Sections were mounted and imaged with the Olympus Slide View. Picrosirius red staining was quantified using ImageJ.

### Mouse tissue extraction and dissociation

Mice were euthanized in a CO_2_ chamber and cervical dislocation was used as a secondary validation of death, as recommended by the IACUC. For liver extraction, perfusion was performed by injecting 5 mL of DPBS (Corning) into the vena cava with a syringe, and the portal vein was cut to allow fluid to drain. Each extracted liver, kidney, or 2 to 3 lungs pooled together from mice in the same group were briefly washed in PBS, then transferred into an individual gentleMACS C Tube (Miltenyi Biotec) containing 5 mL of appropriate dissociation buffer and minced with scissors. For dissociation of liver and kidney, the dissociation buffer contained 10 mM HEPES, 137 mM NaCl, 1 mg/mL collagenase I (Sigma-Aldrich), 1% BSA, and 0.04 mg/mL DNase I. For lungs, a dissociation buffer containing 10 mM HEPES, 137 mM NaCl, 2 mg/mL collagenase I (Sigma-Aldrich), 1% BSA, and 0.04 mg/mL DNase I was used. The gentleMACS C tubes were then attached to gentleMACS Octo Dissociator with Heaters (Miltenyi Biotec). For lung, the m_lung_01_01 program was run three times, followed by a 30-minute incubation at 37°C on the gentleMACS Octo Dissociator, and 5 seconds of the program m_lung_02_01. For the kidney, m_liver_02_02 program was run twice consecutively, and incubated for 30 minutes at 37°C, during which the m_liver_02_02 program was run twice consecutively halfway through incubation. For liver, the m_liver_01_02 program was run twice consecutively, and incubated for 30 minutes at 37°C, during which the m_liver_01_02 program was run twice consecutively halfway through incubation. Cell suspensions were then passed through a 40-μM cell strainer (Corning) and processed with RBC lysis buffer (Zymo Research). For liver, debris removal solution (Miltenyi Biotec) was used following the cell strainer step and before RBC lysis following the manufacturer’s instructions.

### Rotarod and open field

Mice were trained 3 times 2-3 days before measurement. The following setup was used: ramp, initial speed 5 rpm, final speed 40 rpm, RAMP 120 seconds. During training, mice were returned to the rotarod after falling to remain for the total duration. For the measurement, the time of fall was recorded. Measurements were repeated 3 times and reported as an average for each mouse. For open field (Noldus EthoVision), mice were acclimated for at least one hour in the testing room. Mice were placed in the EthoVision field of view and spontaneous activity was recorded for 10 minutes and analyzed using Noldus Ethovision software.

### Flow cytometry and fluorescence-activated cell sorting

Single-cell suspensions of IMR-90, MEFs, 3T3, mouse lung, kidney, spleen, or liver were stained with 7-AAD (1:100 dilution) and anti-Lamp1 antibody (1:100 dilution, human, clone H4A3, BioLegend, mouse, REA777, Miltenyi Biotec) in FACS buffer (PBS with 2% FBS or PBS with 0.5% BSA and EDTA) for 30 minutes at 4°C, protected from light. After staining, cells were washed with FACS buffer and then analyzed on MACSQuant Analyzer (Miltenyi Biotec) and/or fluorescence-activated cell sorting on MACSQuant Tyto Sorter (Miltenyi Biotec). To test LAMP1 binding in physiological conditions, trypsinized and washed cells were incubated at 37°C for 15 minutes with the required antibodies.

### Immunofluorescence

SEN and NS cells were washed once in PBS and fixed in 4% PFA for 15 minutes at room temperature. After washing the PFA with PBS 3 times, 0.2% Triton X-100 was added for 15 minutes and then washed with PBS 3 times. Blocking was done using 5% goat serum overnight at 4C. Primary antibodies (1:500) were prepared in 5% goat serum in TBST and incubated overnight at 4C. Secondary antibodies (1:1000) and Hoechst (1:2000) were added for one hour at room temperature in the dark. After washing with PBS, representative images were taken with an EVOS microscope.

### Real-time cytotoxicity assay (xCELLigence)

ADC toxicity to senescent IMR-90 was measured using the xCELLigence platform (Agilent). Cytotoxicity was normalized to positive untreated controls, IgG controls, and negative full-lysis controls. 50 nM of Anti-Mouse IgG Fc-Duocarmycin Antibody with Cleavable Linker (MORADEC, AM-102-DD) was added to the wells for two hours after one hour of anti-LAMP1 incubation (26 nM). Impedance measurements to assess cell viability were performed every 15 minutes for 24-48 hours. All experiments were performed in triplicates in at least three independent experiments.

### Quantitative RT-PCR

RNA was extracted from mouse tissue cells with Qiagen RNeasy Plus mini kit following manufacturer’s instructions. cDNA was synthesized with PrimeScript RT reagent Kit (Takara Bio) following the manufacturer’s instructions. Real-time PCR was done using AzuraQuant Green Fast qPCR Mix LoRox (RealTimePrimers) on QuantStudio3 (Applied Biosystems, Thermo Fisher) in 96-well plates, and gene expression levels were calculated using the 2^-ΔΔCt^ method. Unless otherwise specified, Actb was used as the housekeeping control. Mouse *Hprt* was used as the housekeeping control in the bleomycin-treated mice lungs.

PCR primers (primer 1; primer 2):

Lmnb1: CAACTGACCTCATCTGGAAGAAC; TGAAGACTGTGCTTCTCTGAGC

Glb1: CTTCCCACTGAACACTGAGGC; TTGGCACGAACAAGGTCTTTT

p16: CCGAACTCTTTCGGTCGTAC; AGTTCGAATCTGCACCGTAGT

p21: TGTCGCTGTCTTGCACTCTG; GACCAATCTGCGCTTGGAGT

Tgf-b: TGATACGCCTGAGTGGCTGTCT; CACAAGAGCAGTGAGCGCTGAA

Hprt: GCTGACCTGCTGGATTACAT; TTGGGGCTGTACTGCTTAAC

Gapdh: CTGGAGAAACCTGCCAAGTA; TGTTGCTGTAGCCGTATTCA

Actb: AAGAGCTATGAGCTGCCTGA; TACGGATGTCAACGTCACAC

### Pathway analysis and correlation analysis

To analyze the proteomics screen of the plasma membrane of senescent cells we used the online tool Enrichr (https://maayanlab.cloud/Enrichr/enrich). The list of 636 genes was obtained from Marin et al., 2023 and a hard threshold was set for upregulation in at least three out of seven senescence induction models (Marin et al., 2023). Enrichr presented as KEGG 2021 Human, BioPlanet 2019 or GO Cellular Component 2023, as indicated in the figure legends. Web interface tool Correlation AnalyzeR was used to study the correlation between LAMP1 and genes of interest, with a ‘Basic’ corGSEA annotation (https://gccri.bishop-lab.uthSENa.edu/shiny/correlation-analyzer/).

### Single-cell RNA Sequencing analysis

The scRNA-Seq data from the ‘Human Muscle Ageing Cell Atlas (HLMA)’ was analyzed using the online tool (https://db.cngb.org/cdcp/hlma/rnaseq/). CDKN1A- and LAMP1-expressing cells were filtered based on > 2 expression to identify cell clusters expressing p21-high LAMP1-high (Lai et al., 2024).

### Statistical methods

Graph Pad Prism 10 was used to perform statistical analysis. Images were processed using ImageJ. Flow cytometry data was analyzed using FlowLogic. Unless otherwise specified, all data are presented as mean ± S.E.M. Unless otherwise specified, all experiments are representative of at least 3 independent experiments. Comparisons between groups were done using an unpaired t-test or ordinary one-way ANOVA. Statistical parameters are defined in the figure legends.

## Author contributions

G.M.L. and A. Sharma wrote the manuscript. M.R., and A.B., extensively reviewed the manuscript. G.M.L., M.Q., and A. Sharma designed the study. G.M.L. and M.Q. performed most experiments and analyzed results. A.B., and A. Shankar performed experiments and analyzed results. All authors participated in discussion and editing the manuscript. All authors have read and agreed to the published version of the manuscript.

## Acknowledgments

This work was supported by the SENS Research Foundation (SRF), Lifespan Research Institute, and VitaDAO. We would like to acknowledge the donors of SENS Research Foundation, Lifespan Research Institute and VitaDAO for sponsoring this work. We would like to acknowledge Yafei Hou for his contributions to discussions and generation of preliminary data for this manuscript. We would like to acknowledge Esmeralda Jimenez for her contributions caring for the animals, conducting daily wellness checks and maintenance, and for providing valuable insights on animal welfare and experimental procedures.

## Data availability statement

Data is available upon reasonable request.

